# Single-cell landscape of immunological responses in COVID-19 patients

**DOI:** 10.1101/2020.07.23.217703

**Authors:** Ji-Yuan Zhang, Xiang-Ming Wang, Xudong Xing, Zhe Xu, Chao Zhang, Jin-Wen Song, Xing Fan, Peng Xia, Jun-Liang Fu, Si-Yu Wang, Ruo-Nan Xu, Xiao-Peng Dai, Lei Shi, Lei Huang, Tian-Jun Jiang, Ming Shi, Yuxia Zhang, Alimuddin Zumla, Markus Maeurer, Fan Bai, Fu-Sheng Wang

## Abstract

In COVID-19 caused by SARS-CoV-2 infection, the relationship between disease severity and the host immune response is not fully understood. Here we performed single-cell RNA sequencing in peripheral blood samples of five healthy donors and 13 COVID-19 patients including moderate, severe and convalescent cases. Through determining the transcriptional profiles of immune cells, coupled with assembled T cell receptor and B cell receptor sequences, we analyzed the functional properties of immune cells. Most cell types in COVID-19 patients showed a strong interferon-alpha response, and an overall acute inflammatory response. Moreover, intensive expansion of highly cytotoxic effector T cell subsets, such as CD4^+^ Effector-GNLY (Granulysin), CD8^+^ Effector-GNLY and NKT CD160, was associated with convalescence in moderate patients. In severe patients, the immune landscape featured a deranged interferon response, profound immune exhaustion with skewed T cell receptor repertoire and broad T cell expansion. These findings illustrate the dynamic nature of immune responses during the disease progression.

---

The coronavirus disease 2019 (COVID-19) caused by severe acute respiratory syndrome coronavirus 2 (SARS-CoV-2) infection causes a spectrum of illness from mild to severe disease and death^1^. Though SARS-CoV-2 like SARS-CoV uses angiotensin-converting enzyme 2 (ACE2) as its receptor for entry in target cells^2, 3^, the viral shedding pattern is different between the two viruses. The SARS-CoV-2 viral load is detectable during the presymptomatic stage^4-6^ and peaks soon after disease onset^7-9^, which is significantly earlier than that of SARS-CoV^10^. These factors contribute to the high contagious nature of SARS-CoV-2 and its rapid spread, leading to the global pandemic of COVID-19. Both SARS-CoV-2 infection and viral infection-mediated-immune responses can directly and/or indirectly damage cells in the respiratory tract of COVID-19 patients^11, 12^. The majority of the patients exhibit mild to moderate symptoms, up to 15% progress to severe pneumonia and approximately 5% eventually develop acute respiratory distress syndrome (ARDS) and/or multiple organ failure^11^. Higher fatality rates have been observed in elderly individuals with co-morbidities and those who are immunocompromised^13-16^.

Since there are no effective drugs or vaccines available at this time against SARS-CoV-2, there is an urgent need to better understand the host immune response during disease in order to better devise prognostic and diagnostic markers, and to design appropriate therapeutic interventions for patients with severe disease presentation.

Viral infection and the antiviral host immune response interact *in vivo* and shape disease severity as well as clinical outcomes, especially during acute viral infection. Therefore, the immunopathology of COVID-19 has received much attention. Immune responses in a COVID-19 patient with moderate disease presentation^17^, show that a robust cellular and humoral immune response can be elicited upon acute SARS-CoV-2 infection. However, it remained unknown how the uncontrolled innate and impaired adaptive immune responses were associated with pulmonary tissue damage. COVID-19 patients with severe disease presentation showed pronounced lymphopenia and elevation of serum pro-inflammatory cytokines^18, 19^. We recently reported a fatal case where significant interstitial lymphocytic infiltrates in both lung tissues and overactivation of T cells in peripheral blood were observed^20^. More recently, inflammatory FCN1^+^ macrophages were found to replace FABP4^+^ macrophages in the bronchoalveolar lavage fluid from severe SARS-CoV-2 infected patients, whereas highly expanded and functional competent tissue resident clonal CD8^+^ T cells were observed in moderate patients with SARS-CoV-2 infection^21^. These observations have revealed possible immunopathogenic mechanisms underlying COVID-19 progression at the first glance. However, a global characterization of the antiviral or pathogenic immune responses in different clinical settings is still lacking.

Here, we implemented single-cell RNA sequencing (scRNA-seq) to obtain an unbiased and comprehensive visualization of the immunological responses in peripheral blood mononuclear cells (PBMCs) from COVID-19 patients with moderate to severe symptoms. Our study depicts a high-resolution transcriptomic landscape of blood immune cells during the disease progression of COVID-19, which will facilitate a better understanding of the protective and pathogenic immune responses of the disease.

## Results

### Single-cell transcriptional profiling of peripheral immune cells

To characterize the immunological features of patients with COVID-19, we performed droplet-based scRNA-seq (10X Genomics) to study the transcriptomic profiles of peripheral blood mononuclear cells (PBMCs) from 13 patients and five healthy donors (HD: n=5) as controls (Fig. 1a). The 13 COVID-19 patients were classified into three clinical conditions: moderate (Moderate: n=7), severe (Severe: n=4) and convalescent (Conv: n=6 of whom four were paired with moderate cases) (Fig. 1a, b, Table 1 and Supplementary Fig. 1). The clinical characteristics and laboratory findings of enrolled patients are detailed in Table 1. Single cell T cell receptor (TCR) and B cell receptor (BCR) sequencing were also performed for each subject. After the unified single cell analysis pipeline (see Methods), ∼0.6 billion unique transcripts were obtained from 122,542 cells from PBMCs of all samples. Among these cells, 22,711 cells (18.5%) were from the HD condition, 37,901 cells (30.9%) were from the Moderate condition, 24,640 cells (20.1%) were from the Severe condition, and 37,290 cells (30.4%) were from the Conv condition. All high-quality cells were integrated into an unbatched and comparable dataset and subjected to principal component analysis after correction for read depth and mitochondrial read counts (Supplementary Fig. 2a, b). Using graph-based clustering (uniform manifold approximation and projection, UMAP), we captured the transcriptomes of 14 major cell types or subtypes according to the expression of canonical gene markers (Fig. 1c-e and Supplementary Fig. 2a, b). These cells included naive state T cells (Naive T: *CD3*^+^*CCR7*^+^), activated state T cells (Activated T: *CD3*^+^*PRF1*^+^), MAIT cells (*SLC4A10*^+^*TRAV1-2*^+^), γδT cells (*TRGV9*^+^*TRDV2*^+^), proliferative T cells (Pro T: *CD3*^+^*MKI67*^+^), natural killer (NK) cells (*KLRF1*^+^), B cells (*MS4A1*^+^), plasma B cells (*MZB1*^+^), CD14^+^ monocytes (CD14^+^ Mono: *LYZ*^+^*CD14*^+^), CD16^+^ monocytes (CD16^+^ Mono: *LYZ*^+^*FCGR3A*^+^), monocyte-derived dendritic cells (Mono DCs: *CD1C*^+^), plasmacytoid dendritic cells (pDCs: *LILRA4*^+^), platelets (*PPBP*^+^) and hemopoietic stem cells (HSCs: *CYTL1*^+^*GATA2*^+^). As such, we clearly defined the composition of cell subpopulations in peripheral blood.

**Table 1.**
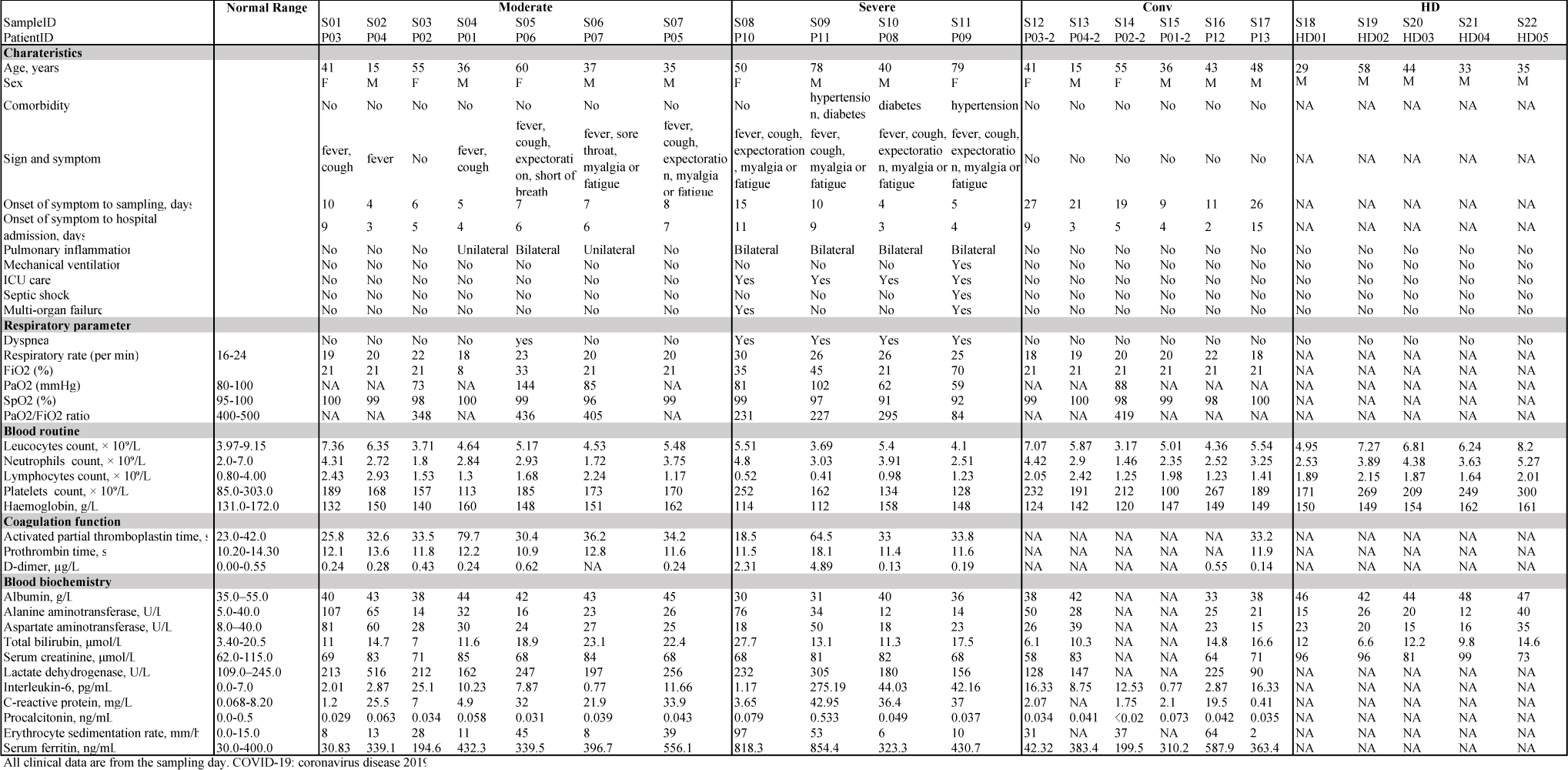
Characteristics and laboratory findings of healthy donors and enrolled patients infected with SARS-CoV-2 in the study.

**Fig. 1.**
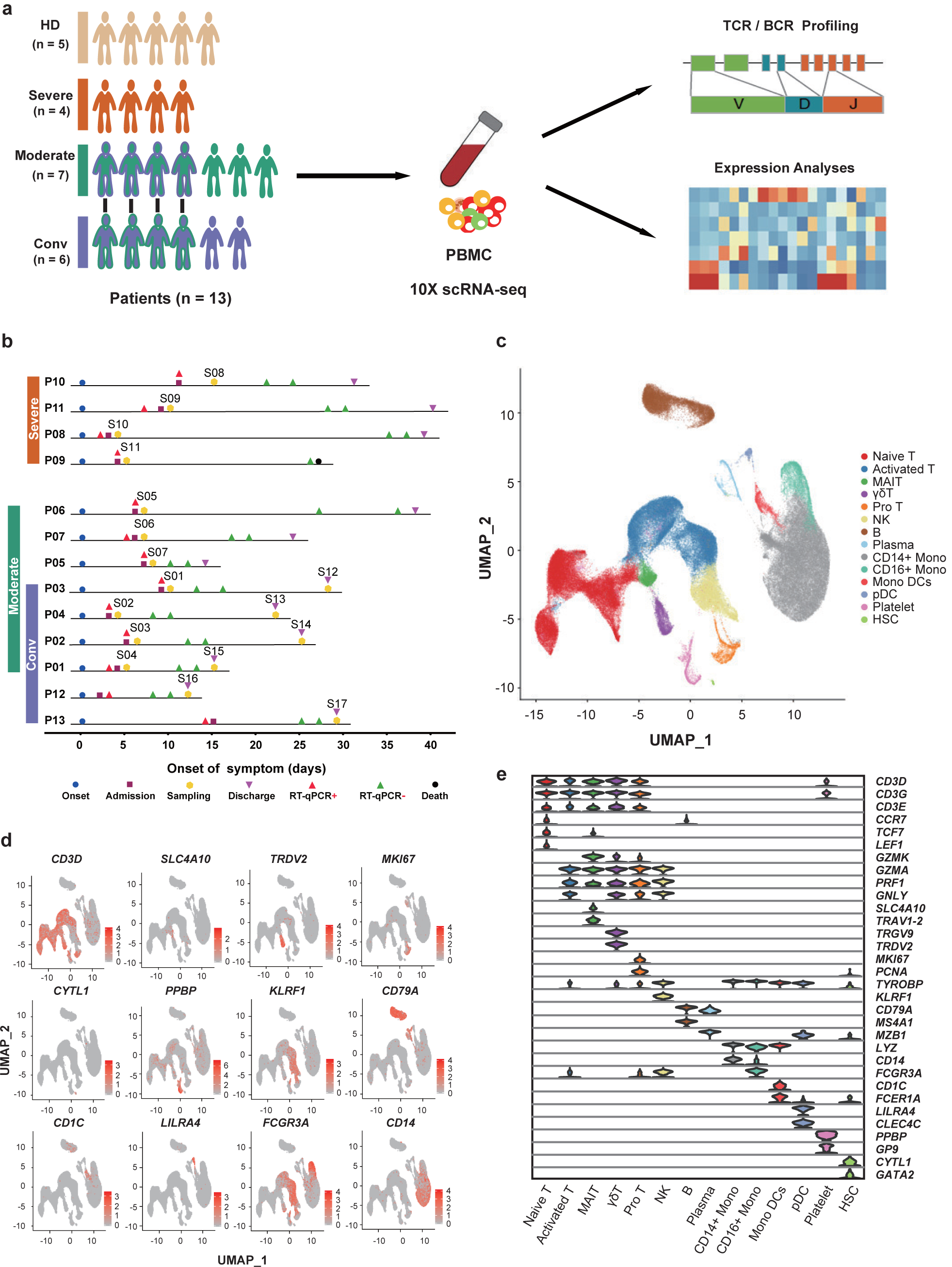
Study design and single-cell transcriptional profiling of PBMCs from healthy donors and COVID-19 patients. **a**, A schematic showing the overall study design. Single-cell RNA sequencing was applied to peripheral blood mononuclear cells across four conditions, and the output data were used for TCR and BCR profiling and expression analyses. **b**, Timeline of the course of disease for 13 SARS-CoV-2 infected patients enrolled in our study. RT-qPCR indicates polymerase chain reaction test for SARS-CoV-2 nucleic acids. RT-qPCR positive indicate nasopharyngeal or sputum samples positive for SARS-CoV-2 nucleic acids. The color bars in the most left representing conditions with the same color in (**a**). **c**, Cellular populations identified. The UMAP projection of 122,542 single cells from HD (n=5), Moderate (n=7), Severe (n=4) and Conv (n=6) samples, showing the formation of 14 clusters with the respective labels. Each dot corresponds to a single cell, colored according to cell types. **d**, Canonical cell markers were used to label clusters by cell identity as represented in the UMAP plot. Colored according to expression levels and legend labeled in log scale. **e**, Violin plots showing the expression distribution of selected canonical cell markers in the 14 clusters. Row representing selected marker genes and column representing clusters with the same color in (**c**).

### Differences in cell compositions across disease conditions

To reveal the differences in cell compositions across three conditions (Moderate, Severe, and Conv), and to compare with that in HD, we calculated the relative percentage of the 14 major cell types in the PBMCs of each individual on the basis of scRNA-seq data (Fig. 2a-d). The relative percentage of activated T cluster peaked in moderate patients and did not return to the normal level, even at convalescence. Of note, the relative abundance of naive T cells, MAIT cells, and Mono DCs decreased with the disease severity, and these populations later restored in Conv patients (Fig. 2d). In contrast, the relative percentage of Pro T cells, plasma B cells, CD14^+^ Mono, and platelets increased with disease severity and later declined in Conv patients (Fig. 2d). The massive increase of CD14^+^ Mono in patients with severe disease was in accordance with a recent study demonstrating that inflammatory monocytes, induced by pathogenic T cells, incite the inflammatory storm in COVID-19^22^.

**Fig. 2.**
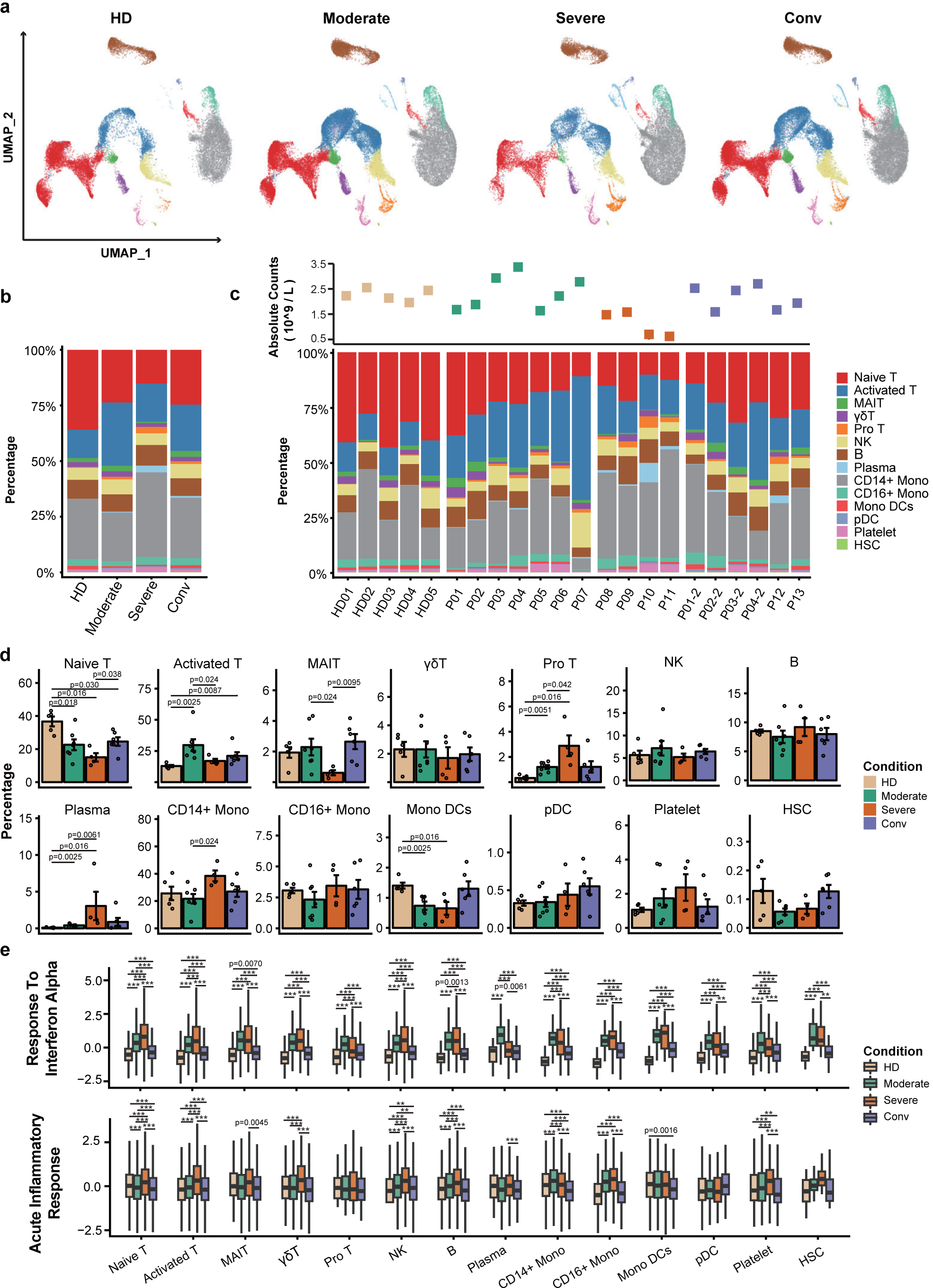
Differences of cell compositions across disease conditions. **a**, UMAP projection of the HD, Moderate, Severe and Conv condition. Each dot corresponds to a single cell, colored according to cell type. **b**, Average proportion of each cell type derived from HD (n=5), Moderate (n=7), Severe (n=4) and Conv (n=6) samples. **c**, The top dot plot shows the sum of the absolute counts of lymphocytes and monocytes in the PBMCs of each sample. The bottom bar plot shows the cell compositions at a single sample level. **d**, Condition preference of each cluster. Y axis: average percent of samples across four conditions. Conditions are shown in different colors. Each bar plot represents one cell cluster. Error bars represent ± s.e.m. for five healthy donors and 13 patients. All differences with *P* < 0.05 are indicated; two-sided unpaired Mann-Whitney *U* test. **e**, Box plots of the expression levels of two GO biological process terms across clusters derived from HD (n=5), Moderate (n=7), Severe (n=4) and Conv (n=6) samples. Conditions are shown in different colors. Horizontal lines represent median values, with whiskers extending to the farthest data point within a maximum of 1.5 × interquartile range. All differences with *P* < 0.01 are indicated. \*\**P* < 0.001; \*\*\**P* < 0.0001; two-sided unpaired Dunn’s (Bonferroni) test.

Next, to investigate the antiviral and pathogenic immune responses during SARS-CoV-2 infection, we evaluated the expression levels of two important pathways (Gene Ontology biological process terms: Response to interferon alpha, and Acute inflammatory response) in major cell types across four conditions. We found that the response to interferon alpha was uniformly and significantly up-regulated in all major cell types from the PBMCs of COVID-19 patients and showed the highest value in almost every major cell type in severe patients, with the exception of plasma B cells, in which the interferon alpha response was greatest in moderate patients (Fig. 2e). In addition, with the exception of Pro T cells, the acute inflammatory response showed consistent and significant differences across conditions in the selected cell types. Several cell types showed trends in the acute inflammatory response that roughly corresponded with disease severity, including activated T, γδT, NK and CD16^+^ mono (Fig. 2e). Moreover, the plasma levels of Type I IFN, IFN-γ and other inflammatory cytokines displayed the highest levels in severe patients (Supplementary Fig. 2c). These results suggest a strong overall pro-inflammatory response in COVID-19 patients (Fig. 2e).

### Strong interferon responses were observed in innate immune cells

To further investigate the transcriptomic changes of innate immune cells (Fig. 3a, b) after SARS-CoV-2 infection, we compared the expression patterns of the Moderate or Severe condition with that of the HD condition in CD14^+^ and CD16^+^ monocytes. We found that the significantly differentially expressed genes (DEGs) were involved in interferon responses, myeloid leukocyte activation, cytokine production and NF-κB signaling pathway in COVID-19 patients (Fig. 3c, d). In addition, more DEGs in monocytes from the Severe condition were enriched in molecule metabolic and catabolic processes, as well as cytokine secretion (Supplementary Fig. 3a). For NK cells, similar to monocytes, DEGs associated with interferon responses, cytokine production, NF-κB signaling pathway and leukocyte cytotoxicity were significantly enriched in COVID-19 patients (Fig. 3e, f), suggesting a consistent response by innate immune cells to SARS-CoV-2 infection. Furthermore, in comparison with moderate patients, the DEGs of the NK cells of the Severe condition, such as *ITGB2, CCL5* and *CXCR2*, were more closely related to migration associated processes (Supplementary Fig. 3b, c).

**Fig. 3.**
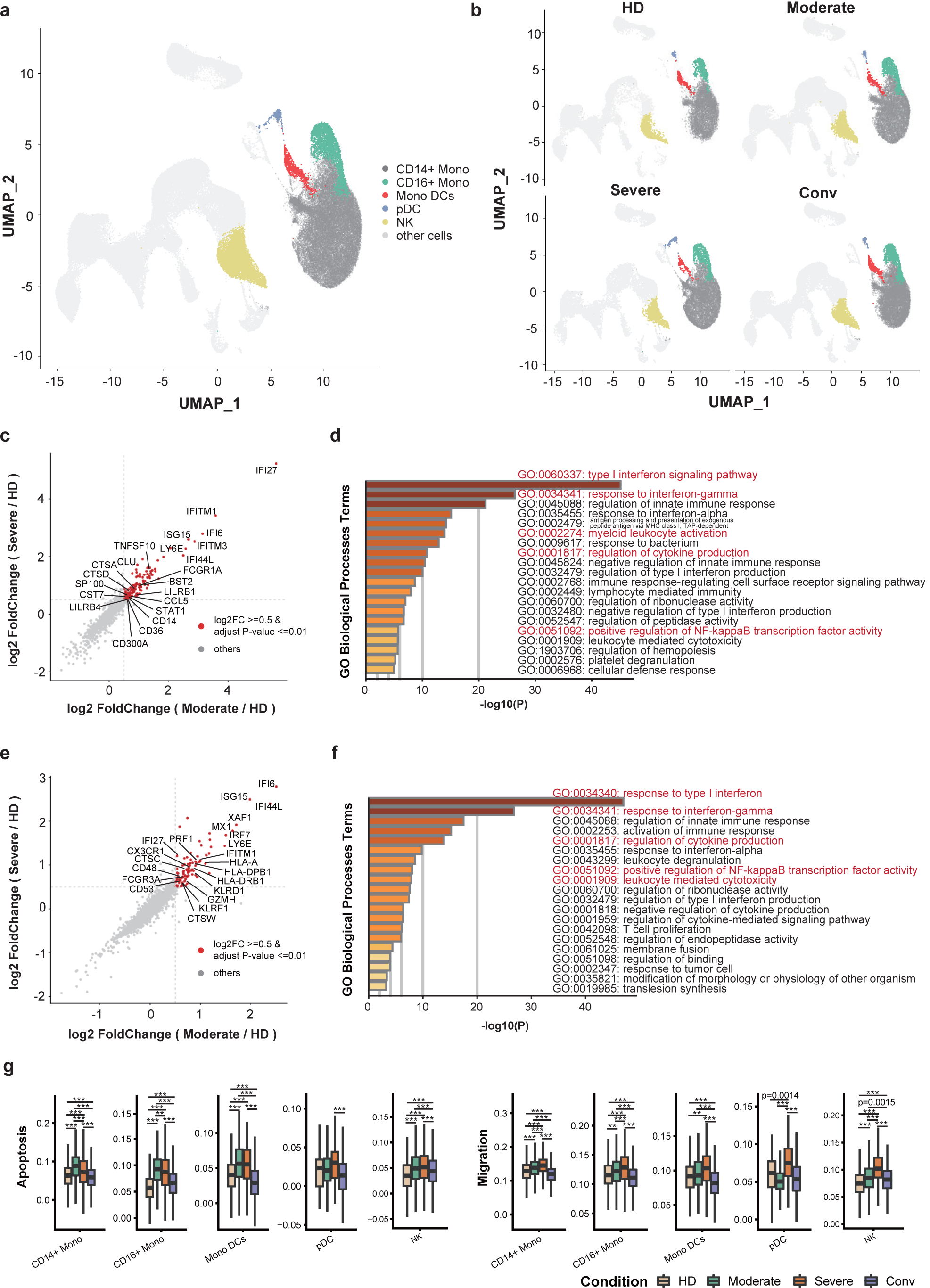
Characterization of innate immune cells in individuals across four conditions. **a**, UMAP projection of monocytes (39,276) and NK cells (7,479). Each dot corresponds to a single cell, colored according to cell type. **b**, UMAP projection of the HD, Moderate, Severe and Conv conditions. **c**, Scatter plot showing DEGs in the monocytes of moderate (n=7) or severe patients (n=4) in comparison with those of HDs (n=5). Each red dot denotes an individual gene with Benjamini-Hochberg adjusted *P* value (two-sided unpaired Mann-Whitney *U* test) ≤ 0.01 and average log2 fold change ≥ 0.5 for the Moderate/HD and Severe/HD comparisons. Example genes are labeled with a gene name. **d**, Gene enrichment analyses of the DEGs colored in red in (**c**). GO terms are labeled with name and id, and sorted by -log10 (*P*) value. A darker color indicates a smaller p-value. The top 20 enriched GO terms are shown. Interesting terms are labeled in red color. **e**, Similar to (**c**), but for NK cells. **f**, Similar to (**d**), but for NK cells. **g**, Box plots of the expression levels of two GO biological process terms in monocytes and NK cells across clusters derived from HD (n=5), Moderate (n=7), Severe (n=4) and Conv (n=6) samples. Conditions are shown in different colors. Horizontal lines represent median values, with whiskers extending to the farthest data point within a maximum of 1.5 × interquartile range. All differences with *P* < 0.01 are indicated. \*\**P* < 0.001; \*\*\**P* < 0.0001; two-sided unpaired Dunn’s (Bonferroni) test.

In line with the DEG enrichment results, we found that monocytes and NK cells both showed significantly up-regulated IFN and acute inflammatory responses after SARS-CoV-2 infection, especially in severe patients (Fig 2e). Levels of cellular apoptosis and migration were also up-regulated in both monocytes and NK cells compared to the HD condition (Fig. 3g). Unlike the comparable apoptosis levels in monocytes and NK cells, the innate immune cells in severe patients were more prone to migration than those in moderate patients (Fig. 3g and Supplementary Fig. 3c). These results suggest that most innate immune cell types in COVID-19 patients show strong interferon responses.

### Features of T cell subsets in COVID-19 patients

To characterize changes in individual T cell subsets among individuals across four conditions, we sub-clustered T cells from PBMCs and obtained 12 subsets according to the expression and distribution of canonical T cell markers (Fig. 4a, b): 6 subtypes of CD4^+^ T cells (*CD3E*^+^*CD4*^+^), three subtypes of CD8^+^ T cells (*CD3E*^+^*CD8A*^+^) and 3 subtypes of NKT cells (*CD3E*^+^*CD4*^−^*CD8A*^−^*TYROBP*^+^).

Of the six subtypes of CD4^+^ T cell clusters, in addition to naive CD4^+^ T (CD4^+^ Naive: *CCR7*^*+*^*SELL*^*+*^), memory CD4^+^ T (CD4^+^ Memory: *S100A4*^*+*^*GPR183*^*+*^), effector memory CD4^+^ T (CD4^+^ Effector Memory: *S100A4*^*+*^*GPR183*^*+*^*GZMA*^*+*^) and regulatory T (Treg: *FOXP3*^*+*^*IL2RA*^*+*^) subtypes, we defined two effector CD4^+^ T subtypes, CD4^+^ Effector-GZMK and CD4^+^ Effector-GNLY. The CD4^+^ Effector-GNLY cluster was characterized by high expression of genes associated with cytotoxicity, including *NKG7, GZMA, GZMB, GZMH* and *GNLY*, while the CD4^+^ Effector-GZMK cluster showed a relatively high expression level of the *GZMK* gene, but low expression of other cytotoxic genes (Fig. 4b and Supplementary Fig. 4a, b). Furthermore, CD4^+^ Effector-GNLY cells showed high expression of *TBX21*, implying that they were Type I T helper (T_H_1) liked cells (Supplementary Fig. 4c). In contrast, CD4^+^ Effector-GZMK and CD4^+^ Effector Memory harbored Type II T helper (T_H_2) liked features with high expression of *GATA3* (Supplementary Fig. 4c). The three subtypes of CD8^+^ T cell clusters included naive CD8^+^ T subset (CD8^+^ Naive: *CCR7*^*+*^*SELL*^*+*^), and two effector CD8^+^ T subsets (CD8^+^ Effector-GZMK and CD8^+^ Effector-GNLY), which both had high expression levels of *GZMA* and *NKG7*. In detail, CD8^+^ Effector-GZMK uniquely expressed *GZMK*, whereas CD8^+^ Effector-GNLY showed relatively high expression levels of *GZMB/H* and *GNLY* (Fig. 4b and Supplementary Fig. 4a, b). The three subsets of NKT cell clusters were defined as naive NKT cells (NKT Naive: *CCR7*^*+*^*SELL*^*+*^), CD56^+^ NKT cells (NKT CD56) and CD160^+^ NKT cells (NKT CD160) (Fig. 4a, b).

**Fig. 4.**
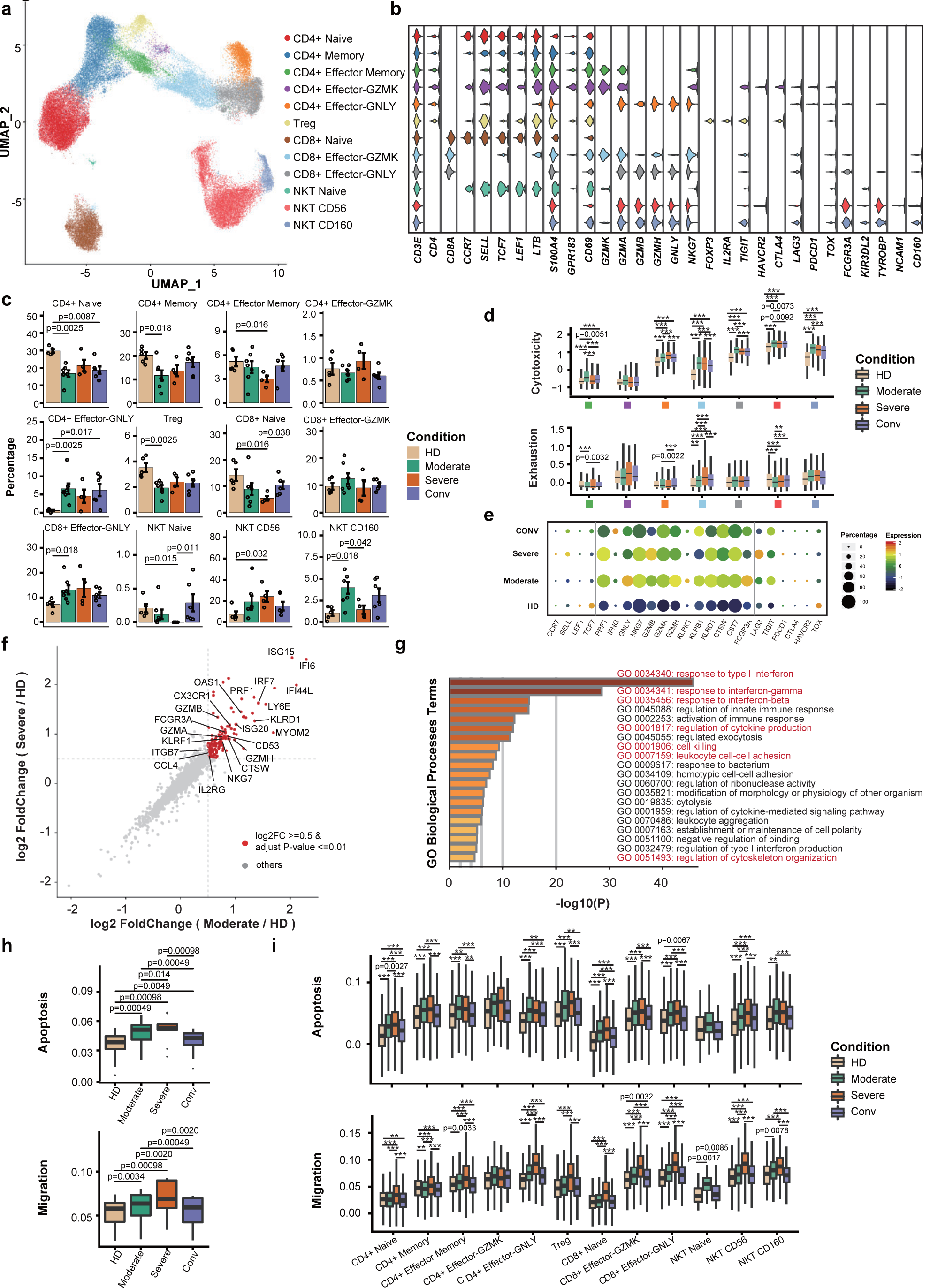
Immunological features of T cell subsets. **a**, UMAP projection of 55,655 T cells. Each dot corresponds to a single cell, colored according to cell type. **b**, Violin plots showing the expression distribution of canonical cell markers in 12 T cell subsets. **c**, Condition preference of each cluster. Error bars represent ± s.e.m. for five healthy donors and 13 patients. All differences with *P* < 0.05 are indicated; two-sided unpaired Mann-Whitney *U* test. **d**, Box plots of the cytotoxicity and exhaustion scores across different clusters and conditions. The squares in the X axis indicate subsets corresponding to subsets in (**a**). Horizontal lines represent median values, with whiskers extending to the farthest data point within a maximum of 1.5 × interquartile range. All differences with *P* < 0.01 are indicated. \*\**P* < 0.001; \*\*\**P* < 0.0001; two-sided unpaired Dunn’s (Bonferroni) test. **e**, Dot plot showing the expression levels of well-defined cytotoxic and exhaustion-related genes in NKT CD160 cells across four conditions. **f**, DEGs of moderate (n=7) or severe patients (n=4) in comparison with HDs (n=5). Each red dot denotes an individual gene with Benjamini-Hochberg adjusted *P* value (two-sided unpaired Mann-Whitney *U* test) ≤ 0.01 and average log2 fold change ≥ 0.5 in the Moderate/HD and Severe/HD comparisons. Example genes are labeled with the gene name. **g**, Gene enrichment analyses of DEGs colored in red in (**f**). Interesting GO terms are labeled in red. **h**, Box plots of the median cell scores for each cluster for two GO biological process terms across HD (n=5), Moderate (n=7), Severe (n=4) and Conv (n=6) samples. Horizontal lines represent median values, with whiskers extending to the farthest data point within a maximum of 1.5 × interquartile range. All differences with *P* < 0.05 are indicated; two-sided paired Mann-Whitney *U* test. **i**, The expression levels of two GO biological process terms across clusters derived from HD (n=5), Moderate (n=7), Severe (n=4) and Conv (n=6) samples. Horizontal lines represent median values, with whiskers extending to the farthest data point within a maximum of 1.5 × interquartile range. All differences with *P* < 0.01 are indicated. \*\**P* < 0.001; \*\*\**P* < 0.0001; two-sided unpaired Dunn’s (Bonferroni) test.

To gain insights into features in T cell subsets, we evaluated the distribution of each cluster across four conditions (Fig. 4c and Supplementary Fig. 4d, e). Notably, the proportions of naive state T subsets including CD4^+^ Naive, CD4^+^ Memory, CD4^+^ Effector Memory, Treg, CD8^+^ Naive and NKT Naive subsets, decreased in COVID-19 patients in comparison with HDs (Fig. 4c and Supplementary Fig. 4d). Even in the Conv condition, the proportions of CD4^+^ Naive, CD8^+^ Naive and Treg clusters did not restore to the levels of HDs (Fig. 4c and Supplementary Fig. 4d). In contrast, the proportions of active state T subsets, including CD4^+^ Effector-GNLY, CD8^+^ Effector-GNLY, NKT CD56 and NKT CD160 subsets, increased in COVID-19 patients, and these cytotoxic subsets were present in high proportions even in Conv patients (Fig. 4c). Of particular interest, the CD4^+^ Effector-GNLY subset was almost absent in HDs but highly enriched in moderate, severe and Conv patients. In addition, the abundance of the NKT CD160 subset was significantly reduced in severe patients as compared to moderate patients.

We then evaluated the cytotoxicity and exhaustion scores of different effector state T subsets across four conditions (Fig. 4d and Supplementary Fig. 4f). The CD4^+^ Effector-GNLY, CD8^+^ Effector-GNLY, NKT CD56 and NKT CD160 subsets showed higher cytotoxicity scores than those of the other subsets. Within these highly cytotoxic clusters, healthy donors all had the lowest cytotoxicity scores, whereas the Moderate condition showed the highest cytotoxic state, with the exception of the CD4^+^ Effector-GNLY subset (Fig. 4d, e and Supplementary Fig. 4f). Meanwhile, the CD4^+^ Effector-GZMK, CD8^+^ Effector-GZMK and NKT CD160 clusters showed higher exhaustion scores than those of the other subsets. Within these highly exhausted subsets, healthy donors all had the lowest exhaustion scores, while severe patients showed the most exhausted state (Fig. 4d, e and Supplementary Fig. 4f), in agreement with previous functional studies which examined CD8^+^ T cells from severe patients and found highly exhausted status and functional impairment^23^.

To further investigate differential transcriptomic changes in T cells after SARS-CoV-2 infection, we compared the expression profiles of effector T cells (excluding CD4^+^ Naive, CD4^+^ Memory, CD8^+^ Naive and NKT Naive clusters) between the Moderate or Severe and HD conditions. We observed that DEGs up-regulated in COVID-19 patients were involved in processes including interferon responses, cytokine production, cell killing, leukocyte cell-cell adhesion and cytoskeleton organization (Fig. 4f, g and Supplementary Fig. 4i). In addition, using an apoptosis and migration scoring system, we observed that T cells in severe patients likely underwent migration and apoptosis (Fig. 4h, i and Supplementary Fig. 4g, h). Significant activation of cell death and migration pathways in the PBMCs of severe patients suggests that cell death and lymphocytes migration may be associated with lymphopenia, a common phenomenon observed in severe COVID-19 patients^18, 19, 24^.

### Clonal expansion in T cells and preferred usage of V(D)J genes in COVID-19 patients

Next, to gain insight into the clonal relationship among individual T cells and usage of V(D)J genes across four conditions, we reconstructed TCR sequences from the TCR sequencing. Briefly, there were more than 70% of cells in all subsets with matched TCR information except for the three NKT subsets (Fig. 5a, b). First, compared to the HDs, clonal expansion was obvious in COVID-19 patients and patients in convalescence (Fig. 5c-e). The extent of clonal expansion in the Moderate and Conv conditions was higher than that of the Severe condition. Meanwhile, large clonal expansions (clonal size > 100) were absent in the Severe condition (Fig. 5e), indicating that severe patients might lack efficient clonal expansion of effector T cells. We observed different degrees of clonal expansion among T cell subsets (Fig. 5c, d). Effector T cell subsets CD4^+^ Effector-GNLY, CD8^+^ Effector-GZMK and CD8^+^ Effector-GNLY showed high proportions of clonal cells (Fig. 5a, d and Supplementary Fig. 5a) and contained high proportions of inter-cluster clonal cells (Fig. 5f), suggesting that effector T cells underwent dynamic state transitions (Fig. 5a, f).

**Fig. 5.**
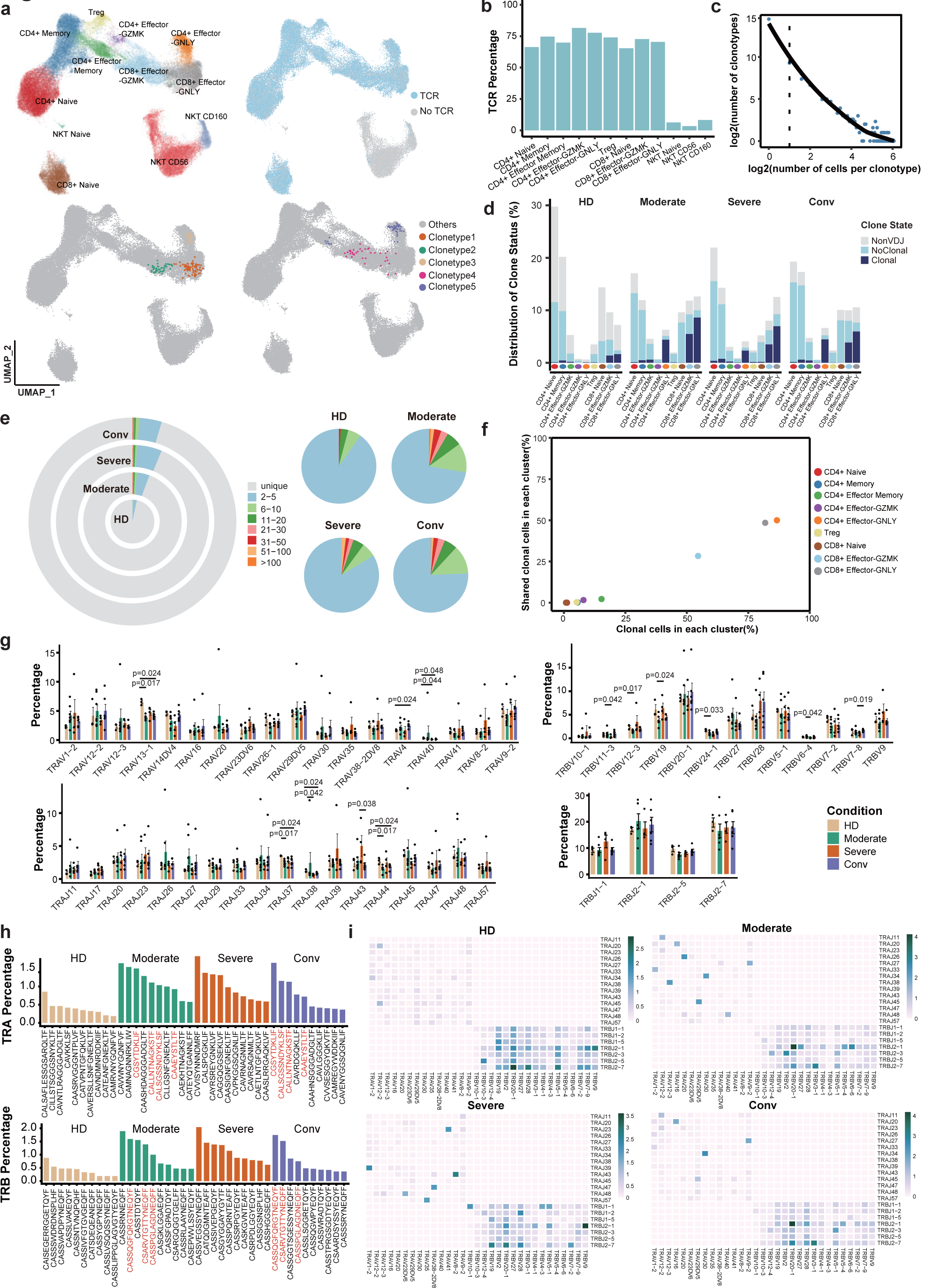
Expanded TCR clones and selective usage of V(D)J genes. **a**, UMAP of T cells derived from PBMCs. Clusters are denoted by color labeled with inferred cell types (top left), TCR detection (top right), selected TCR clones belonging to the same clusters (bottom left) and different clusters (bottom right). **b**, Bar plots showing the percentage of TCR detection in each T cell cluster. **c**, The association between the number of T cell clones and the number of cells per clonotype. The dashed line separates non-clonal and clonal cells. LOESS fitting is labeled as the solid line showing negative correlation between the two axes. **d**, The distribution of the clone state of T cells in each cluster. **e**, The clonal status percentage of T cells (left) and percentage of different levels of clonal T cells (right) across four conditions. **f**, Comparison between the fraction of clonal cells in each subset (X-axis) and percentage of cells with TCRs shared across clusters (Y-axis). **g**, Usage of some TRAV (top left), TRAJ (bottom left), TRBV (top right) and TRBJ (bottom right) genes across four conditions. Error bars represent ± s.e.m. for five healthy donors and 13 patients. All differences with *P* < 0.05 are indicated; two-sided unpaired Mann-Whitney *U* test. **h**, The top ten CDR3 usages are shown. Each bar is colored by condition identity. Shared CDR3 sequences are in a red font. **i**, TRA/B rearrangements differences across four conditions. The colors indicate the usage percentage of specific V-J gene pairs.

To study the dynamics and genes preference of TCRs in COVID-19 patients and HDs, we compared the usage of V(D)J genes across four conditions (Fig. 5g-i and Supplementary Fig. 5b). The top 10 complementarity determining region 3 (CDR3) sequences were different across four conditions (Fig. 5h). The Moderate and Conv conditions shared some CDR3 sequences because four samples from these conditions were paired. The usage percentage of the top 10 CDR3 sequences in the HD condition was lower and more balanced compared to those of the other three conditions. Of note, we discovered a different usage of V(D)J genes with decreased diversity in COVID-19 patients, which was more pronounced in TRA genes (Fig. 5i). We also identified over-representation of *TRAJ39* and *TRAJ43* in severe patients compared to moderate and Conv patients (Fig. 5g). The preferred TRBJ gene in severe patients was *TRBJ1-1*, whereas *TRBJ2-1* was preferred in moderate and Conv patients (Fig. 5i). The selective usage of V(D)J genes indicates that different immunodominant epitopes may drive the molecular composition of T cell responses and may be associated with SARS-CoV-2 specific infection.

### Features of B cell subsets in COVID-19 patients

To trace the dynamic changes of different B subtypes, we sub-clustered B cells into six subsets according to the expression and distribution of canonical B cell markers (Fig. 6a, b and Supplementary Fig. 6a). We identified one naive B subset (Naive B; *MS4A1*^*+*^*IGHD*^*+*^), one memory B subset (Memory B; *MS4A1*^*+*^*CD27*^*+*^), one intermediate transition memory B subset (Intermediate Memory B; *IGHD*^*+*^*CD27*^*+*^), one germinal center B subset (Germinal Center B; *MS4A1*^*+*^*NEIL1*^*+*^) and two plasma subsets, Plasma B (Plasma B; *MZB1*^*+*^*CD38*^*+*^) and dividing plasma B (Dividing Plasma B; *MZB1*^*+*^*CD38*^*+*^*MKI67*^*+*^).

**Fig. 6.**
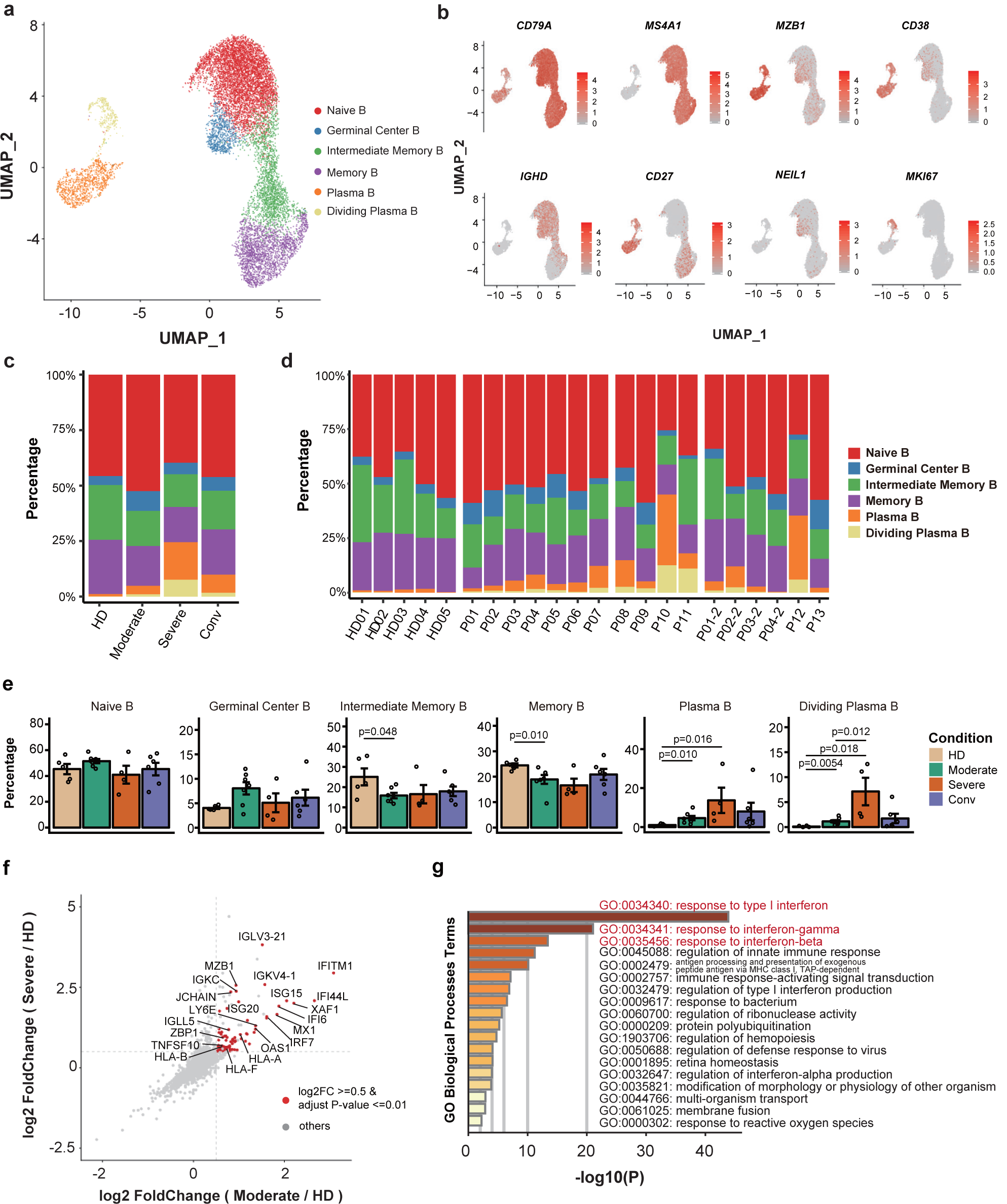
Immunological features of B cell subsets. **a**, UMAP projection of 11,377 B cells. Each dot corresponds to a single cell, colored according to cell type. **b**, Canonical cell markers were used to label clusters by cell identity as represented in the UMAP plot. Colored according to expression level and legend labeled in log scale. **c**, Average proportion of each B cell subtype derived from HD (n=5), Moderate (n=7), Severe (n=4) and Conv (n=6) samples. **d**, Bar plot showing B cell compositions at the single sample level. **e**, Condition preference of each cluster. Y axis: average percentage of samples across four conditions. Conditions are shown in different colors. Each bar plot represents one cell cluster. Error bars represent ± s.e.m. for five healthy donors and 13 patients. All differences with *P* < 0.05 are indicated; two-sided unpaired Mann-Whitney *U* test. **f**, Scatter plot showing DEGs in moderate (n=7) or severe (n=4) patients in comparison with those of HDs (n=5). Each red dot denotes an individual gene with Benjamini-Hochberg adjusted *P* value (two-sided unpaired Mann-Whitney *U* test) ≤ 0.01 and average log2 fold change ≥ 0.5 in the Moderate/HD and Severe/HD comparisons. Example genes are labeled with the gene name. **g**, Gene enrichment analyses of DEGs colored in red in (**f**). GO terms are labeled with name and id and sorted by -log10 (*P*) values. A darker color indicates a smaller p-value. The top 20 enriched GO terms are shown. Interesting terms are labeled in red.

Notably, the proportions of active state B subsets, including Germinal Center B, Plasma B and Dividing Plasma B subsets, increased in COVID-19 patients in comparison with those of HDs. In contrast, the proportion of Memory B cells decreased in COVID-19 patients compared to that of HDs (Fig. 6c-e).

To further investigate differential transcriptomic changes in B cells after SARS-CoV-2 infection, we compared the expression profiles of B/Plasma cells of the Moderate or Severe condition to those of the HD condition. DEGs that were most significantly enriched in COVID-19 patients were involved in genes associated with the interferon response (Fig 6f, g and Supplementary Fig. 6c). Moreover, DEGs in severe patients were associated with protein synthesis, maturation and transport related biological processes (Supplementary Fig. 6b). These results reveal the transcriptomic features of B cell subsets in COVID-19 patients.

### Expanded B cells and specific rearrangements of V(D)J genes in severe patients

We also reconstructed B cell receptor (BCR) sequences from BCR sequencing and analyzed the state of B cell-receptor (BCR) clonal expansion. Briefly, the detection percentage of BCRs was more than 75% in each cluster (Fig. 7a, b). We found that B cells from severe patients showed obvious clonal expansions (Fig. 7c and Supplementary Fig. 6d) than other three conditions, indicating that B cell activity and humoral immune responses were strongly activated in severe patients, reminiscent of previous observation that higher antibody titers are associated with worse clinical outcomes^25-27^. This raises the concern that, pathogen-directed antibodies can promote disease pathology, resulting in antibody-dependent enhancement (ADE) similar to that observed in SARS^28^.

**Fig. 7.**
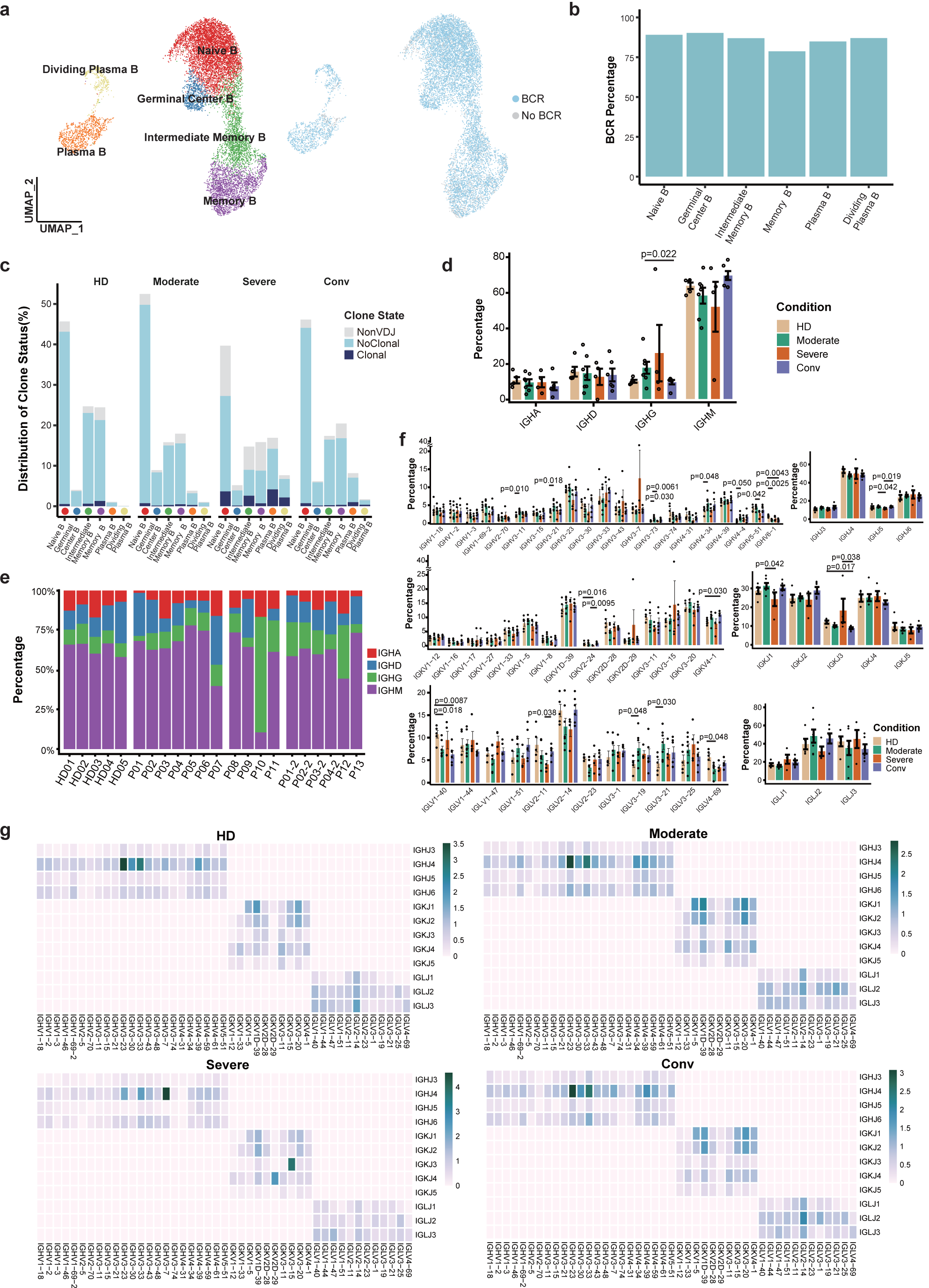
Expanded BCR clones and selective usage of V(D)J genes. **a**, UMAP of B cells derived from PBMCs. Clusters are denoted by color and labeled with inferred cell types (left). The UMAP of B cells is also colored based on BCR detection (right). **b**, Bar plot showing the percentage of BCR detection in each B cell cluster. **c**, Bar plot showing the distribution of the clone state of B cells in each cluster. **d**, Bar plot showing the percentages of IGHA, IGHD, IGHG and IGHM in each condition, with error bars representing ± s.e.m. for five healthy donors and 13 patients. All differences with *P* < 0.05 are indicated; two-sided unpaired Mann-Whitney *U* test. **e**, Bar plots showing the percentages of IGHA, IGHD, IGHG and IGHM at the single sample level. **f**, Usage of some IGHV (top left), IGHJ (top right), IGKV (middle left), IGKJ (middle fight), IGLV (bottom left) and IGLJ (bottom right) genes across the HD, Moderate, Severe and Conv conditions. Conditions are shown in different colors. Error bars represent ± s.e.m. for five healthy donors and 13 patients. All differences with *P* < 0.05 are indicated; two-sided unpaired Mann-Whitney *U* test. **g**, Heatmaps showing IGH/K/L rearrangements differences across the HD, Moderate, Severe and Conv conditions. The colors indicate the usage percentages of specific V-J gene pairs.

Next, we evaluated the distribution of IgA, IgD, IgG and IgM (IgE not detected) in each patient at the Moderate, Severe and Conv conditions, respectively. In most patients, IgM was the predominant immunoglobulin (Fig. 7d, e). Compared to HDs, the abundance of IgG increased in COVID-19 patients, while that of IgM levels decreased. In convalescent patients, levels of IgG and IgM returned to levels similar to those of HDs.

To study biased V(D)J rearrangements of the BCR, we compared the usage of V(D)J genes across four conditions (Fig. 7f, g and Supplementary Fig. 6e). We found more specific V(D)J usage in the Severe condition compared with the other three conditions, indicating that B cells might have undergone unique and specific V(D)J rearrangements in severe patients (Fig. 7g). We also discovered comprehensive usage of *IGHJ4* in all HDs and patients (Fig. 7f), but the paired IGHV genes of *IGHJ4* were different in severe patients compared with patients in the other three conditions (Fig. 7g). We observed over-representation of *IGHV3-7* in severe patients (Fig. 7f). Moreover, the top two paired V-J frequencies in severe patients were *IGHV3-7*/*IGHJ4* and *IGKV3-15*/*IGKJ3* (Fig. 7g). Taken together, increased B cell clonality and skewed usage of the IGHV and IGKJ genes in severe patients, suggests that SARS-CoV-2 infection is associated with V(D)J rearrangements in B cells of the host. Notably, selective usage of dominant IGV genes, especially *IGHV3-7* and *IGKV3-15* in severe patients, may facilitate the design of vaccines.

## Discussion

COVID-19 is usually considered as an acute self-limited viral disease^29^, although it remains unknown whether SARS-CoV-2 infection can lead to chronic disease in asymptomatic carriers. Host immune response against acute SARS-CoV-2 infection not only plays an antiviral role, but also leads to simultaneous pathogenic injury of organs and tissues, especially in the lungs of COVID-19 patients, which determines the disease severity, progression and outcome. Studies have reported the characteristics of innate and adaptive immune responses^15, 18, 24^, which have helped us understand the potential pathogenesis of SARS-CoV-2 infection. However, it is difficult to obtain an integrated scenario of the cellular and molecular immune responses upon SARS-CoV-2 infection. To address this issue, here we have profiled the immunological response landscape in COVID-19 patients at single cell resolution, which illustrates the dynamic nature of cellular responses during disease progression and reveals the critical factors responsible for antiviral immunity and pathogenesis in moderate and severe patients.

Our study provides an unbiased visualization of the immunological hallmarks for COVID-19 patients. First, COVID-19 patients showed a concerted and strong interferon alpha response, an overall acute inflammatory response, and an enhanced migration ability, which peaked in patients with severe disease in most major cell types in the PBMCs. Second, broad immune activation was observed in COVID-19 patients, evidenced by increased proportions of activated T, Pro T and plasma B cells, and decreased proportions of naive T and Mono DC compartments. Third, the proportions of active state T clusters were significantly higher in COVID-19 patients and with a preferential enrichment of effector T cell subsets, such as CD4^+^ Effector-GNLY, CD8^+^ Effector-GNLY and NKT CD160 cells in moderate patients and NKT CD56 subset in severe patients. T cells showed higher cytotoxicity and more robust expansion in moderate patients, whereas higher exhaustion levels and less specific clonal expansion were seen in severe patients. Fourth, B cells experienced unique and specific V(D)J rearrangements in severe patients, indicated by an increase of B cell clonality and a skewed use of the IGHV and IGKJ genes. Finally, though most of the clinical parameters recovered to normal range in patients at the early phase of convalescence, the state of the immune system was not fully restored, exemplified by the ratios of naive T and Treg subsets. A long-term follow-up study is needed to investigate how long it takes to achieve full immune recovery in patients with COVID-19.

An effective anti-viral immune response in moderate patients was characterized by moderate and broad activation of innate immune signals as well as expansion of highly cytotoxic effector T cell subsets. The expanded Effector T clusters, including CD4^+^ Effector-GNLY, CD8^+^ Effector-GNLY, NKT CD56 and NKT CD160, share features of high expression of *NKG7, GZMA. GZMB, GZMH* and *GNLY*, and may promote rapid resolution of SARS-CoV-2 infection through their direct cytotoxicity. The CD4^+^ Effector-GNLY cluster resembles classical CD4^+^ cytotoxic T cells^30^. CD4^+^ cytotoxic T cells with MHC class II-restricted cytotoxic activity play an important role in viral infections^31^, autoimmune diseases^32^, and malignancies^33^. Further, it is of great interest to identify immune factors that may predict or prevent progression to severe illness. Notably, the expansion of NKT CD160 cluster in moderate patients is almost absent in the Severe condition. The NKT CD160 cluster refers to a previously described γδ NKT or Vδ1 T cell subset that shows phenotypical and functional similarity to traditional NK cells^34^. Moreover, *FCGR3A* was also among the enriched genes in the NKT CD160 cluster, suggesting that it may mediate the antibody dependent cell-mediated cytotoxicity. Furthermore, Vδ1 T cells are implicated in immune responses to viral infection, particularly cytomegalovirus^35, 36^, Epstein Barr virus^37^, and Vδ1 T cells can also recognize a broad range of cancer cells^38, 39^. As in COVID-19, the preferential expansion of NKT CD160 cells might promote rapid control of the disease through direct cytotoxicity as well as mediating the antibody dependent cell-mediated cytotoxicity effect. Mechanisms underlying the expansion and function of NKT CD160 cells in COVID-19 warrant future studies. It will be valuable to investigate whether Vδ1 T cells could be used in adoptive cellular therapies to curb overt COVID-19 associated tissue and organ damages.

The immunopathogenesis of disease deterioration in severe patients was characterized by a deranged interferon response, profound immune exhaustion with skewed TCR repertoire and broad T cell expansion. Importantly, previous studies on SARS-CoV showed that the virus can harness multiple mechanisms to antagonize interferon responses in host cells^12^. Based on the results of in vitro cell infection, SARS-CoV-2 was recognized as a weak inducer of type I interferons^40, 41^. However, here we observed strong interferon alpha responses in almost all cells types in the PBMCs from severe patients in comparison to moderate patients. Considering that the virus burden peaked soon after disease onset and then decreased gradually^7, 8^, it seems that the systematic interferon alpha signal activation in severe patients might be induced by factors other than the virus alone. Over-activation of interferon pathways may contribute to immune dysfunction and immune injury in severe COVID-19 patients. Particularly, interferon-α2b nebulization was widely applied in SARS-CoV-2 infection, which was developed from treating MERS and SARS^42^. The use of interferon for treatment may need a careful reconsideration and reexamination, especially in severe COVID-19 cases.

There are several limitations in this study. For example, it was very difficult to obtain the immune cells in bronchoalveolar lavage fluid due to biosafety reasons during the outbreak of COVID-19 when we performed this study. Also, the sample size is comparatively small. Therefore, future studies with longitudinal samples from more COVID-19 patients may help to determine the cause-and-effect relationships between immune characteristic of different cell types and disease outcome.

Taken together, this integrated, multi-cellular description in our study lays the foundation for future characterization of the complex, dynamic immune responses to SARS-CoV-2 infection. The transcriptomic data, coupled with detailed TCR- and BCR-based lineage information, can serve as a rich resource for deeper understanding of peripheral lymphocytes in COVID-19 patients and pave the way for rationally designed therapies as well as development of SARS-CoV-2-specific vaccines.

## Acknowledgements

We thank the study participants who provided blood samples, and Chun-Bao Zhou, Jin-Hong Yuan for flow cytometry analysis. Jun Hou provided guidance for safely handling samples from COVID-19 patients. This work was supported by the National Key Research and Development Program of China (grant no. 2020YFC0841900 and 2020YFC0844000) to F-S. W. from the Ministry of Science and Technology of China.

## Author contributions

F-S.W. and F.B. conceived and designed the study; J-Y.Z., X.F., J-W.S., X-P.D. and C.Z. performed the experiments; X-M.W. and X.X. led the bioinformatic analyses; X-M.W., X.X., J-Y.Z., X.F., J-W.S., C.Z., F.B. and F-S.W. wrote the manuscript; J-L.F., P.X., S-Y.W., L.S., Z.X., L.H., and T-J.J. took care of patients and provided the clinical information; R-N.X., M.S., Y.Z., A.Z. and M.M. edited the manuscript and provided comments and feedback. All authors read and approved the final manuscript.

## Methods

### Patient cohort and clinical characteristics

Thirteen COVID-19 patients were admitted at the Fifth Medical Center of PLA General Hospital and enrolled in the study from January 23 to February 15, 2020. The category of three clinical groups (Moderate, Severe and Convalescent (Conv)) were based on Guidelines for diagnosis and treatment of Corona Virus Disease 2019 issued by the National Health Commission of China (7th edition) (http://www.chinacdc.cn/jkzt/crb/zl/szkb_11803/jszl_11815/202003/t20200305_214142.html). Moderate group included non-pneumonia and mild pneumonia cases. Severe group included severe cases who met one of the following criteria: (1) Respiratory distress, respiratory rate ≥30 breaths/min; (2) Pulse oxygen saturation (SpO2) ≤93% without inhalation of oxygen-support at quiet resting state; (3) Arterial partial pressure of oxygen (PaO2) / oxygen concentration (FiO2) ≤300 mmHg; (4) CT image shows there is more than 50% increase of lung infiltrating change within 24 to 48 hours. Critically ill cases who were grouped in the Severe condition in this study generally required mechanical ventilation and exhibited respiratory failure, septic shock, and/or multiple organ dysfunction/failure that required monitoring and treatment in the ICU. One case in Severe group died during the study period. The patients in convalescent group met the discharge criteria as follows: afebrile for more than 3 days, resolution of respiratory symptoms, substantial improvement of chest CT images, and 2 consecutive negative reverse transcription-quantitative polymerase chain reaction (RT-qPCR) tests for viral RNA in respiratory tract swab samples obtained at least 24 hours apart. This study was approved by the ethics committee of the hospital and written informed consents or the telephone call permissions were obtained from each patient or their guardian in the very difficult conditions of early COVID-19 pandemic.

The clinical data and disease course of the 13 patients are shown in Table 1 and Fig. 1b, respectively. Blood sampling for single cell RNA sequencing was usually performed at the time of admission or discharge. CT images for one moderate and one severe case exhibited bilateral ground-glass opacities (Supplementary Fig. 1).

### Quantitative reverse transcription polymerase chain reaction

The throat swab, sputum from the upper respiratory tract and blood were collected from patients at various time-points after hospitalization. Sample collection, processing, and laboratory testing complied with WHO guidance. Viral RNA was extracted from samples using the QIAamp RNA Viral Kit (Qiagen, Heiden, Germany) according to the manufacturer’s instructions. SARS-CoV-2 infected patients were confirmed using a RT-qPCR kit (TaKaRa, Dalian, China) as recommended by the China CDC.

### Preparation of single-cell suspensions

Peripheral venous blood samples were obtained on admission of 13 COVID-19 patients and five healthy donors within 24 hours, placed into vacutainer tubes, and centrifuged at 400 × *g* for 5 min at 4 °C. The time of sampling relative to the onset of symptoms was recorded. Plasma samples were collected and stored at −80 °C until use. For each sample, cell viability exceeded 90%.

### Droplet-based single cell sequencing

Using a Single Cell 5′ Library and Gel Bead Kit (10X Genomics, 1000006) and Chromium Single Cell A Chip Kit (10X Genomics, 120236), the cell suspension (300-600 living cells per microliter as determined by Count Star) was loaded onto a Chromium single cell controller (10X Genomics) to generate single-cell gel beads in the emulsion (GEMs) according to the manufacturer’s protocol. Briefly, single cells were suspended in PBS containing 0.04% BSA. Approximately 10,000 cells were added to each channel, and approximately 6,000 target cells were recovered. Captured cells were lysed and the released RNA was barcoded through reverse transcription in individual GEMs. Reverse transcription was performed on a S1000TM Touch Thermal Cycler (Bio Rad) at 53 °C for 45 min, followed by 85 °C for 5 min, and a hold at 4 °C. cDNA was generated and amplified, after which quality was assessed using an Agilent 4200 (performed by CapitalBio Technology, Beijing). According to the manufacturer’s introduction, single-cell RNA-seq libraries were constructed using a Single Cell 5′ Library and Gel Bead Kit, Single Cell V(D)J Enrichment Kit, Human T Cell (1000005) and a Single Cell V(D)J Enrichment Kit, Human B Cell (1000016). The libraries were sequenced using an Illumina Novaseq6000 sequencer with a paired end 150 bp (PE150) reading strategy (performed by CapitalBio Technology, Beijing).

### Single cell RNA-seq data processing

Raw gene expression matrices were generated for each sample by the Cell Ranger (Version 3.0.2) Pipeline coupled with human reference version GRCh38. The output filtered gene expression matrices were analyzed by R software (Version 3.5.3) with the Seurat^43^ package (Version 3.0.0). In brief, genes expressed at a proportion > 0.1% of the data and cells with > 200 genes detected were selected for further analyses. Low-quality cells were removed if they met the following criteria: 1) < 800 UMIs; 2) < 500 genes; or 3) > 10% UMIs derived from the mitochondrial genome. After removal of low-quality cells, the gene expression matrices were normalized by the *NormalizeData* function, and 2000 features with high cell-to-cell variation were calculated using the *FindVariableFeatures* function. To reduce dimensionality of the datasets, the *RunPCA* function was conducted with default parameters on linear-transformation scaled data generated by the *ScaleData* function. Next, the *ElbowPlot, DimHeatmap* and *JackStrawPlot* functions were used to identify the true dimensionality of each dataset, as recommended by the Seurat developers. Finally, we clustered cells using the *FindNeighbors* and *FindClusters* functions, and performed non-linear dimensional reduction with the *RunUMAP* function with default settings. All details regarding the Seurat analyses performed in this work can be found in the website tutorial (https://satijalab.org/seurat/v3.0/pbmc3k_tutorial.html).

### Multiple dataset integration

To compare cell types and proportions across four conditions, we employed the integration methods described at (https://satijalab.org/seurat/v3.0/integration.html)^44^. The Seurat package (Version 3.0.0) was used to assemble multiple distinct scRNA-seq datasets into an integrated and unbatched dataset. In brief, we identified 2000 features with high cell-to-cell variation as described above. Second, we identified “anchors” between individual datasets with the *FindIntegrationAnchors* function and inputted these “anchors” into the *IntegrateData* function to create a “batch-corrected” expression matrix of all cells, which allowed cells from different datasets to be integrated and analyzed together.

### Sub-clustering of B cells and T cells

B cells and plasma cells were extracted from PBMCs. Next, these major cell types were integrated for further sub-clustering. After integration, genes were scaled to unit variance. Scaling, PCA and clustering were performed as described above. Naive and Activated T cells in PBMCs were also extracted and sub-clustered using the procedure used for B cells.

### Cell type annotation and cluster markers identification

After non-linear dimensional reduction and projection of all cells into two-dimensional space by UMAP, cells clustered together according to common features. The *FindAllMarkers* function in Seurat was used to find markers for each of the identified clusters. Clusters were then classified and annotated based on expressions of canonical markers of particular cell types. Clusters expressing two or more canonical cell-type markers were classified as doublet cells and excluded from further analysis.

### Differential expression genes (DEGs) identification and functional enrichment

Differential gene expression testing was performed using the *FindMarkers* function in Seurat with parameter “test.use=wilcox” by default, and the Benjamini-Hochberg method was used to estimate the false discovery rate (FDR). DEGs were filtered using a minimum log2(fold change) of 0.5 and a maximum FDR value of 0.01. Enrichment analysis for the functions of the DEGs was conducted using the Metascape webtool (www.metascape.org). Gene sets were derived from the GO Biological Process Ontology.

### Defining cell state scores

We used cell scores to evaluate the degree to which individual cells expressed a certain pre-defined expression gene set^45-47^. The cell scores were initially based on the average expression of the genes from the pre-defined gene set in the respective cell. For a given cell i and a gene set j (Gj), the cell score SCj (i) quantifying the relative expression of Gj in cell i as the average relative expression (Er) of the genes in Gj compared to the average relative expression of a control gene set (Gjcont): SCj (i) = average[Er(Gj, i)] – average[Er(Gjcont,i)]. The control gene set was randomly selected based on aggregate expression levels bins which yield a comparable distribution of expression levels and over size to that of the considered gene set. The *AddModuleScore* function in Seurat was used to implement the method with default settings. We used RESPONSE TO INTERFERON ALPHA (GO:0035455), RESPONSE TO INTERFERON BETA (GO:0035456), ACUTE INFLAMMATORY RESPONSE (GO:0002526), APOPTOTIC SIGNALING PATHWAY (GO:0097190), LEUKOCYTE MIGRATION (GO:0050900), 4 well-defined naive markers (*CCR7, TCF7, LEF1* and *SELL*), 12 cytotoxicity associated genes (*PRF1, IFNG, GNLY, NKG7, GZMB, GZMA, GZMH, KLRK1, KLRB1, KLRD1, CTSW* and *CST7*) and 6 well-defined exhaustion markers (*LAG3, TIGIT, PDCD1, CTLA4, HAVCR2* and *TOX*) to define the interferon alpha/beta response, inflammatory response, apoptosis, migration, naive state, cytotoxicity and exhaustion score, respectively.

### TCR and BCR V(D)J sequencing and analysis

Full-length TCR/BCR V(D)J segments were enriched from amplified cDNA from 5′ libraries via PCR amplification using a Chromium Single-Cell V(D)J Enrichment kit according to the manufacturer’s protocol (10X Genomics). Demultiplexing, gene quantification and TCR/BCR clonotype assignment were performed using Cell Ranger (Version 3.0.2) vdj pipeline with GRCh38 as reference. In brief, a TCR/BCR diversity metric, containing clonotype frequency and barcode information, was obtained. For the TCR, only cells with at least one productive TCR alpha chain (TRA) and one productive TCR beta chain (TRB) were kept for further analysis. Each unique TRA(s)-TRB(s) pair was defined as a clonotype. For the BCR, only cells with at least one productive heavy chain (IGH) and one productive light chain (IGK or IGL) were kept for further analysis. Each unique IGH(s)-IGK/IGL(s) pair was defined as a clonotype. If one clonotype was present in at least two cells, cells harboring this clonotype were considered clonal, and the number of cells with such pairs indicated the degree of clonality of the clonotype. Using barcode information, T cells with prevalent TCR clonotypes and B cells with prevalent BCR clonotypes were projected on UMAP plots.

### Plasma cytokines detection

Plasma levels of IFN-γ, IL-6, IL-18, MCP-1, MIP-1α and IP-10 were evaluated by using an Aimplex kit (Beijing Quantobio, China) following manufacturer’s instructions. Type I IFN, represented by IFN-β, was detected by ELISA kit (Cat. EK1236, Multi Sciences, China) according to the manufacture’s protocols. All samples were performed in duplicate.

### Statistics

The statistical tools, methods and threshold for each analysis are explicitly described with the results or detailed in the Figure Legends or Methods sections.

### Data Availability

The raw sequence data reported in this paper have been deposited in the Genome Sequence Archive of the BIG Data Center, Beijing Institute of Genomics (BIG), Chinese Academy of Sciences, under accession number HRA000150 and are publicly accessible at http://bigd.big.ac.cn/gsa-human. The raw source data are available from the corresponding author upon reasonable request.

### Code Availability

Experimental protocols and data analyses pipeline used in our work are following the 10X Genomics and Seurat official website. The analyses steps, functions and parameters used are described in details in Methods section. Custom scripts for analyzing data are available upon reasonable request.

