## Supplemental files for "Single-cell landscape of immunological responses in COVID-19 patients"

### Supplementary Figures

#### Supplementary Figure 1

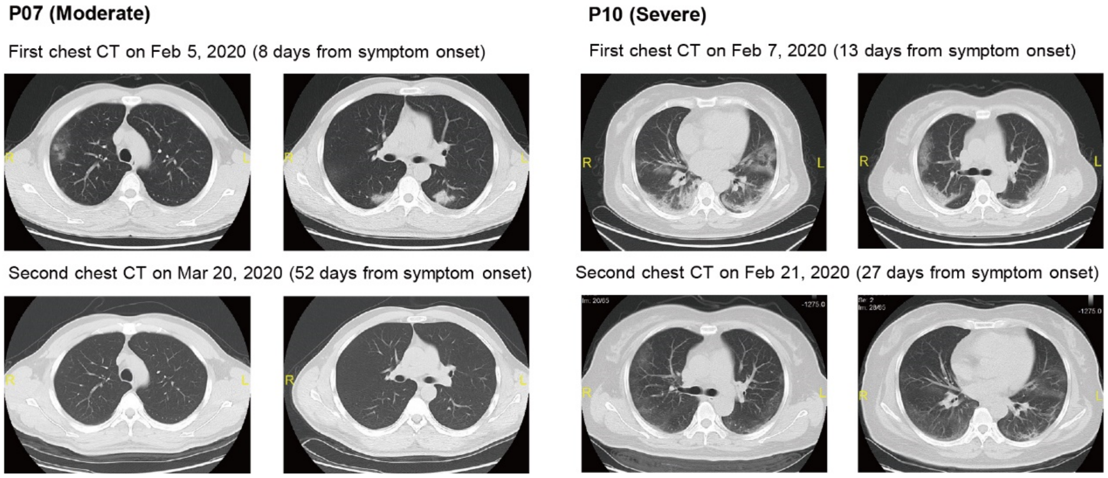


**Supplementary Fig. 1 | Representative chest CT images from moderate and severe cases showing bilateral ground-glass opacity.**

#### Supplementary Figure 2

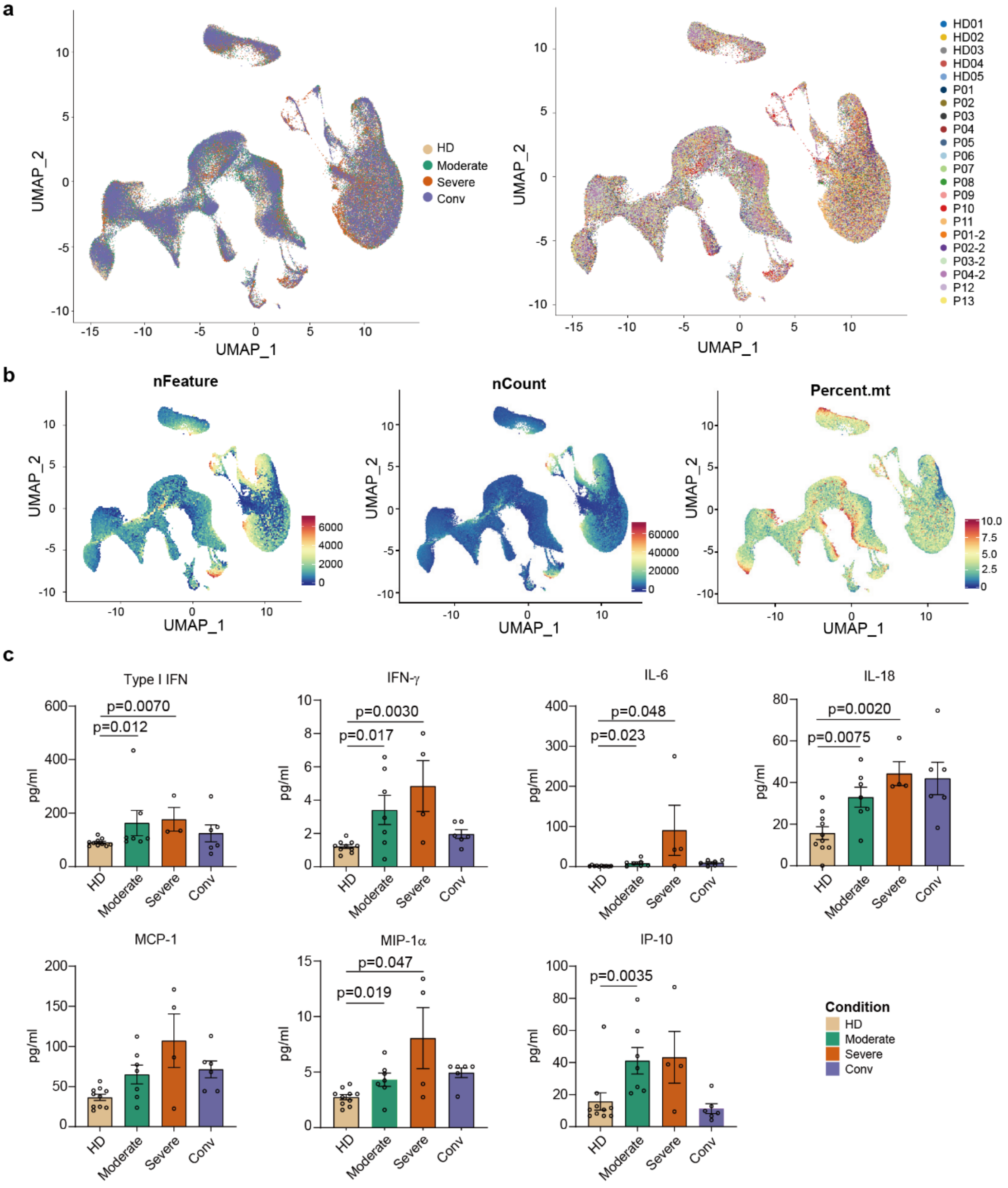


**Supplementary Fig. 2 | Quality of clustering and experiment validation of changes in PBMCs for healthy donors and COVID-19 patients. a,** UMAP of all cells colored by condition identity (left) and sample identity (right). **b,** UMAP of gene counts (left), UMIs (middle) and percentage of mitochondrial genes (right) in all cells. **c,** Plasma interferons and cytokine levels across four conditions: HD (n=10), Moderate (n=7), Severe (n=4) and Conv (n=6) samples. Error bars represent ± s.e.m. for 10 healthy donors and 13 patients. All differences with $P < 0.05$ are indicated; two-sided unpaired Mann-Whitney $U$ test.

#### Supplementary Figure 3

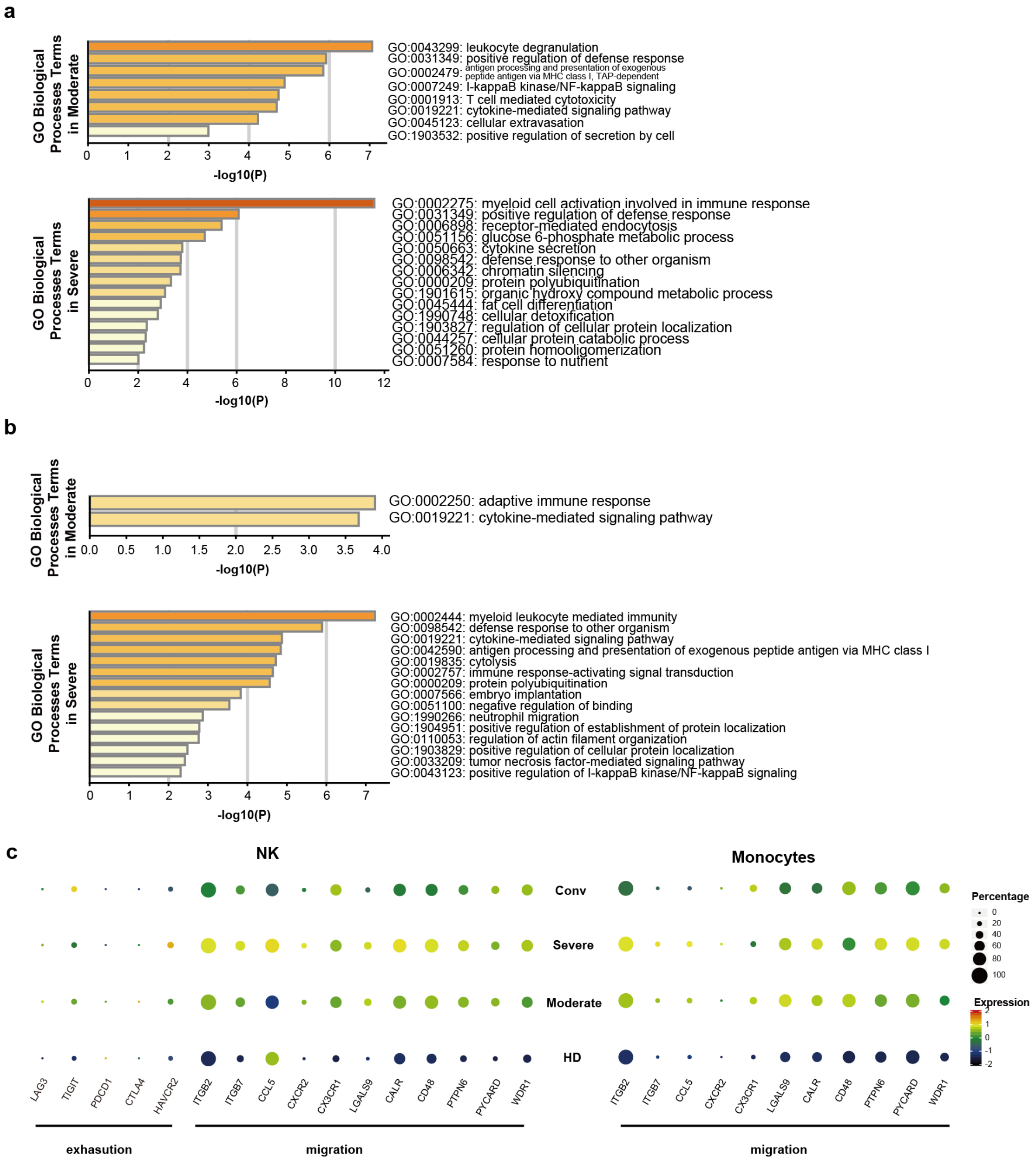


**Supplementary Fig. 3 | Transcriptomic profiling of monocytes and NK cells in Moderate and Severe patients. a,** Gene enrichment analyses of DEGs in monocytes in moderate patients, but not in severe patients, in comparison with healthy donors (top), and genes in severe patients but not in moderate patients comparing to healthy donors (bottom). GO terms were labeled with name and id, and sorted by -log10 (*P*) value. A darker color indicates a smaller p-value. **b,** As in (**a**), but for NK cells. **c**, Dot plot showing expression of genes associated with migration processes in NK cells (left) and monocytes (right) across four conditions.

#### Supplementary Figure 4

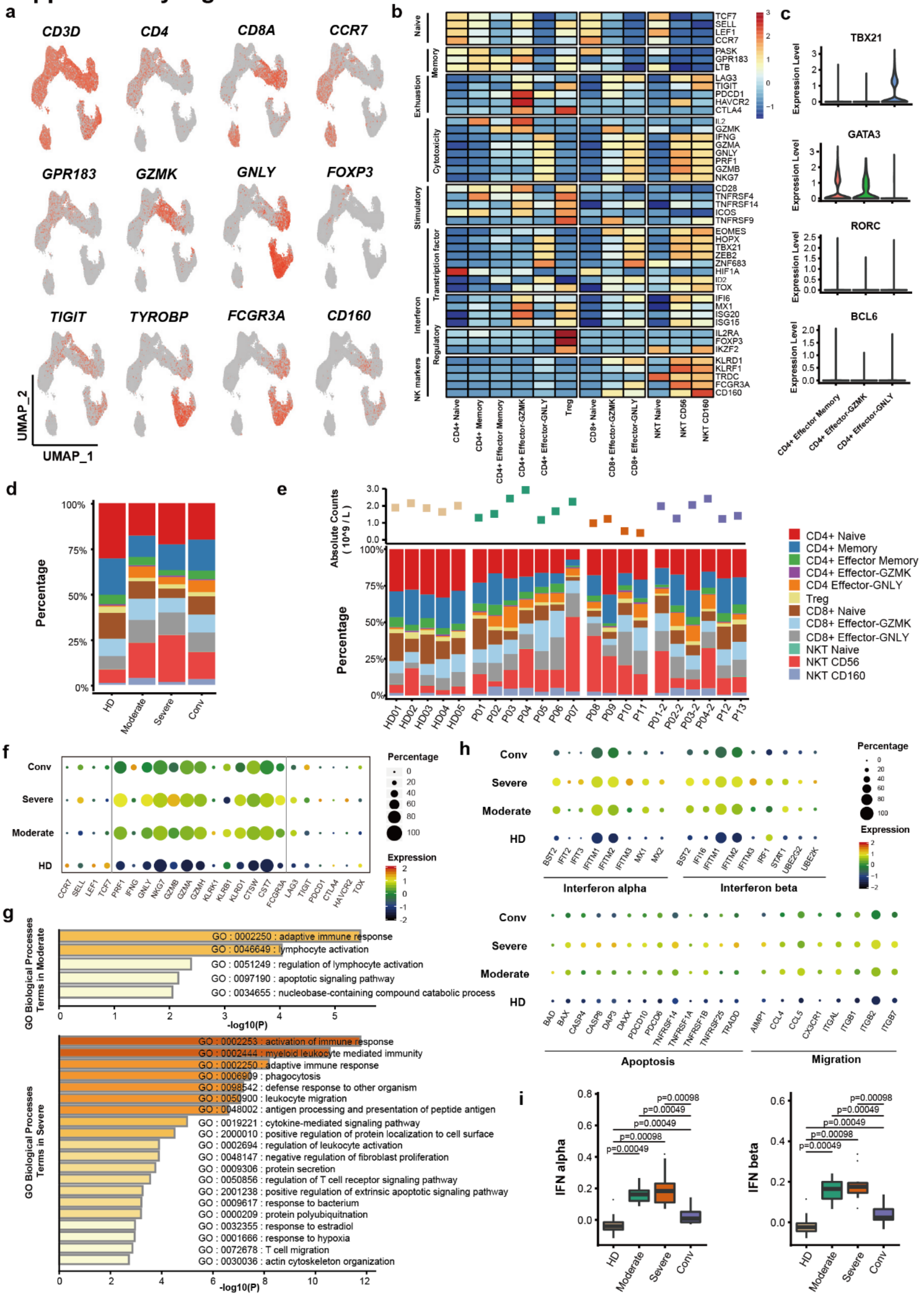


**Supplementary Fig. 4 | Dynamic cell composition and transcriptomic profiles of T cells. a,** Canonical cell markers were used to label clusters by cell identity as represented in the UMAP plot. Clusters are colored according to expression level and the legend is labeled in log scale. **b,** Heatmap showing the z-score normalized mean expression of selected T cell function-associated genes in each cell cluster. **c,** Violin plots showing the expression distribution of selected canonical T helper markers in three effector $CD4^+$ T subset cells. **d,** Average proportion of each T cell subtype derived from HD (n=5), Moderate (n=7), Severe (n=4) and Conv (n=6) samples. **e,** The top plot shows the absolute counts of lymphocytes in PBMC samples. The bottom bar plot shows the T cell compositions at a single-sample level. **f,** Dot plot showing expression of some well-defined naive, cytotoxic and exhausted genes in cells in Fig. 4d across four conditions. **g,** Gene enrichment analyses of DEGs in T cells in moderate patients (top) and in severe patients (bottom) in comparison with healthy donors, respectively. **h,** Dot plot showing expression of some genes associated with interferon alpha and beta response, apoptosis and migration processes across four conditions. **i,** Box plots of the median cell scores for each cluster of interferon alpha and beta response associated genes across HD (n=5), Moderate (n=7), Severe (n=4) and Conv (n=6) samples. Conditions are shown in different colors. Horizontal lines represent median values, with whiskers extending to the farthest data point within a maximum of 1.5 × interquartile range. All differences with $P < 0.05$ are indicated; two-sided paired Mann-Whitney *U* test.

#### Supplementary Figure 5

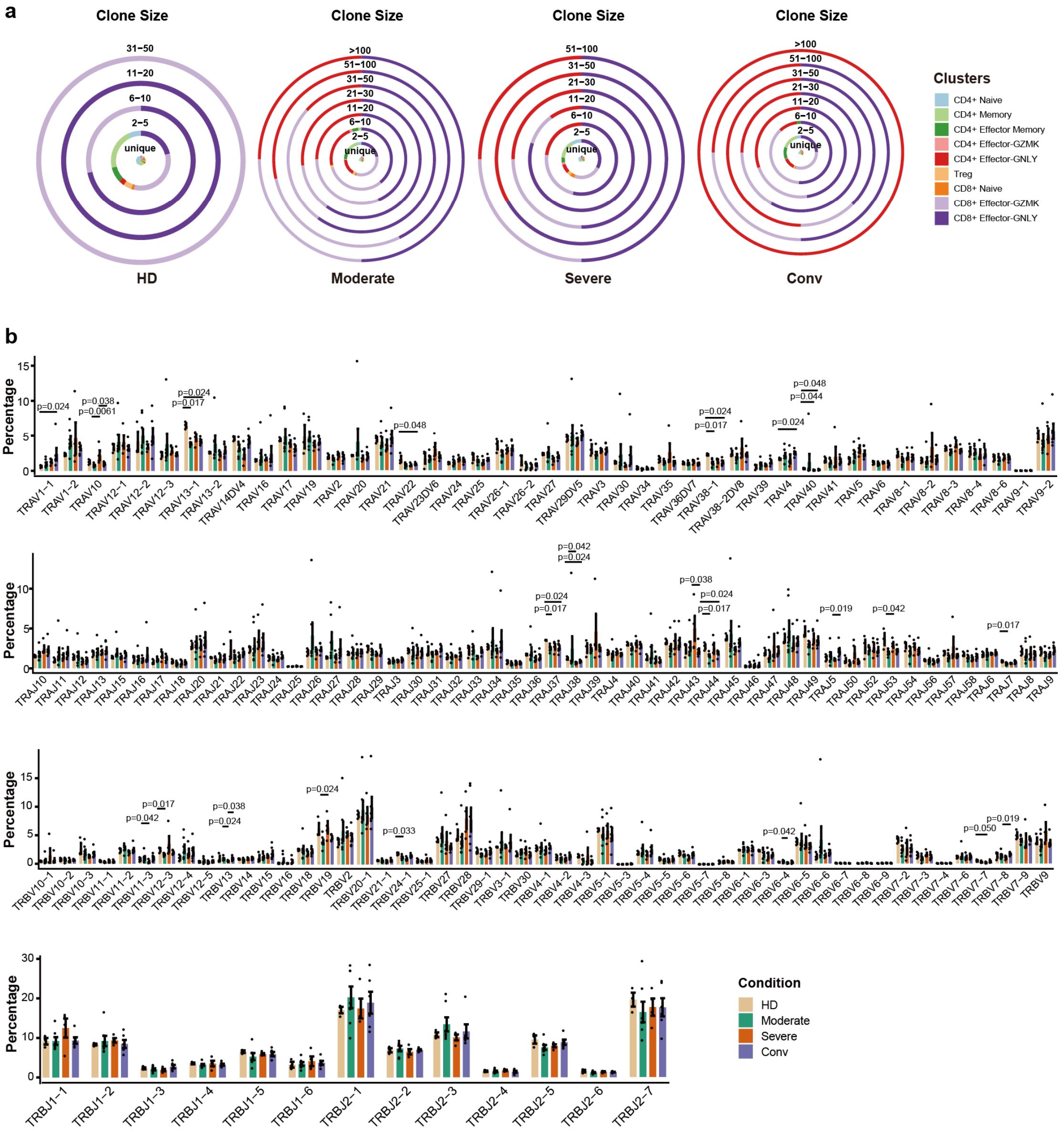


**Supplementary Fig. 5 | Biased expanded TCR clones and usage of V(D)J genes. a,** Pie chart showing the distribution of clonal expanded cells in different clusters across the HD, Moderate, Severe and Conv conditions. **b,** Usage of the TRAV, TRAJ, TRBV and TRBJ genes across the HD, Moderate, Severe and Conv conditions. Conditions are shown in different colors. Error bars represent ± s.e.m. for five healthy donors and 13 patients. All differences with $P < 0.05$ are indicated; two-sided unpaired Mann-Whitney *U* test.

#### Supplementary Figure 6

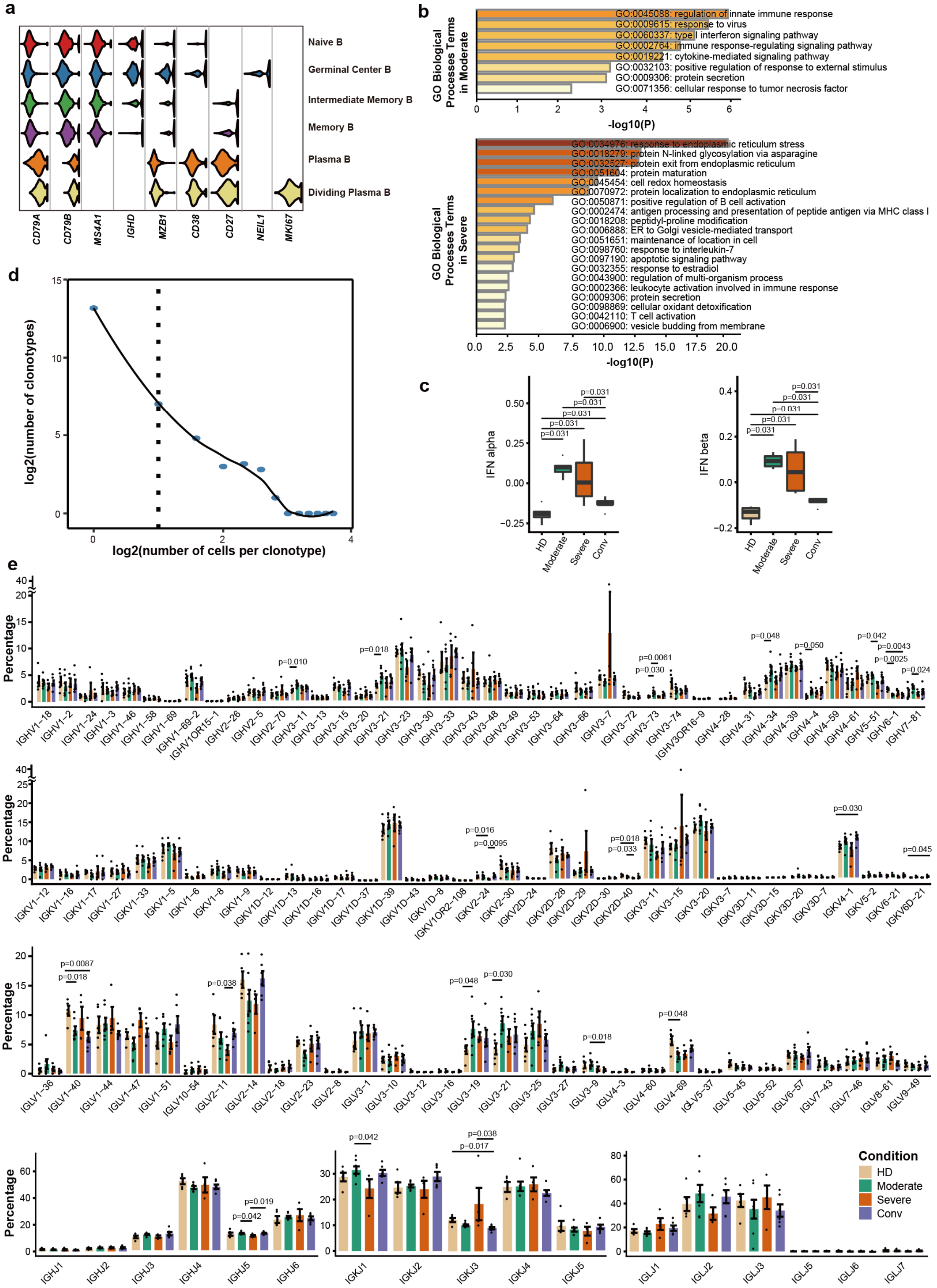


**Supplementary Fig. 6 | Transcriptomic profiling of B cells and biased usage of V(D)J genes.** **a,** Violin plots showing the expression distribution of selected canonical cell markers in six B cell clusters. The columns correspond to selected markers, and the rows correspond to clusters with the same color in Fig. 6a. **b,** Gene enrichment analyses of DEGs in B cells in moderate patients (top) and in severe patients (bottom) in comparison with HDs, respectively. GO terms are labeled with name and id, and sorted by -log10 (*P*) value. **c,** Box plots of median cell scores for each cluster of interferon alpha and beta response associated genes across HD (n=5), Moderate (n=7), Severe (n=4) and Conv (n=6) samples. Conditions are shown in different colors. Horizontal lines represent median values, with whiskers extending to the farthest data point within a maximum of 1.5 × interquartile range. All differences with $P < 0.05$ are indicated; two-sided paired Mann-Whitney *U* test. **d,** Association between the number of B cell clones and the number of cells per clonotype. The dashed line separates non-clonal and clonal cells. LOESS fitting is labeled as the solid line showing negative correlation between the two axes. **e,** Usage of the IGHV/J, IGKV/J and IGLV/J genes across four conditions. Conditions are shown in different colors. Error bars represent ± s.e.m. for five healthy donors and 13 patients. All differences with $P < 0.05$ are indicated; two-sided unpaired Mann-Whitney *U* test.
